# Differential role of cytosolic Hsp70s in longevity assurance and protein quality control

**DOI:** 10.1101/2020.06.25.170670

**Authors:** Rebecca Andersson, Anna Maria Eisele-Bürger, Sarah Hanzén, Katarina Vielfort, David Öling, Frederik Eisele, Gustav Johansson, Tobias Gustafsson, Kristian Kvint, Thomas Nyström

**Affiliations:** Department of microbiology and immunology, Institute of Biomedicine, Sahlgrenska Academy, University of Gothenburg, 403 50 Gothenburg, Sweden; Department of chemistry and molecular biology, Faculty of Science, University of Gothenburg, 403 50 Gothenburg, Sweden

## Abstract

70 kDa heat shock proteins (Hsp70) are essential chaperones of the protein quality control network; vital for cellular fitness and longevity. The four cytosolic Hsp70’s in yeast, Ssa1-4, are thought to be functionally redundant but the absence of Ssa1 and Ssa2 causes a severe reduction in cellular reproduction and accelerates replicative aging. In our efforts to identify which Hsp70 activities are most important for longevity assurance, we systematically investigated the capacity of Ssa4 to carry out the different activities performed by Ssa1/2 by overproducing Ssa4 in cells lacking these Hsp70 chaperones. We found that Ssa4, when overproduced in cells lacking Ssa1/2, rescued growth, mitigated aggregate formation, restored spatial deposition of aggregates into protein inclusions, and promoted protein degradation. In contrast, Ssa4 overproduction in the Hsp70 deficient cells failed to restore the recruitment of the disaggregase Hsp104 to misfolded/aggregated proteins, to fully restore clearance of protein aggregates, and to bring back the formation of the nucleolus-associated aggregation compartment. Exchanging the nucleotide-binding domain of Ssa4 with that of Ssa1 suppressed this ‘defect’ of Ssa4. Interestingly, Ssa4 overproduction extended the short lifespan of *ssa1*Δ *ssa2*Δ mutant cells to a lifespan comparable to, or even longer than, wild type cells, demonstrating that Hsp104-dependent aggregate clearance is not a prerequisite for longevity assurance in yeast.

**AUTHOR SUMMARY:** All organisms have proteins that network together to stabilize and protect the cell throughout its lifetime. One of these types of proteins are the Hsp70s (heat shock protein 70). Hsp70 proteins take part in folding other proteins to their functional form, untangling proteins from aggregates, organize aggregates inside the cell and ensure that damaged proteins are destroyed. In this study, we investigated three closely related Hsp70 proteins in yeast; Ssa1, 2 and 4, in an effort to describe the functional difference of Ssa4 compared to Ssa1 and 2 and to answer the question: What types of cellular stress protection are necessary to reach a normal lifespan? We show that Ssa4 can perform many of the same tasks as Ssa1 and 2, but Ssa4 doesn’t interact in the same manner as Ssa1 and 2 with other types of proteins. This leads to a delay in removing protein aggregates created after heat stress. Ssa4 also cannot ensure that misfolded proteins aggregate correctly inside the nucleus of the cell. However, this turns out not to be necessary for yeast cells to achieve a full lifespan, which shows us that as long as cells can prevent aggregates from forming in the first place, they can reach a full lifespan.

## INTRODUCTION

A major task of cellular Protein Quality Control (PQC) is to prevent protein misfolding and unspecific protein aggregation. Molecular chaperones are central components of such PQC and the heat shock protein 70 (Hsp70) chaperones are among the most conserved members of the PQC system. Originally identified in *Drosophila*, Hsp70 homologs have been found in almost all organisms studied (1). Yeast has 14 genes encoding Hsp70 chaperones; nine cytosolic and five organelle/compartment-specific. These Hsp70s aid in essential housekeeping functions, including *de novo* protein folding, protein translocation, and protein degradation (2).

The general structural domains of Hsp70 chaperones include a 44 kDa amino-terminal ATPase domain (NBD), an 18 kDa substrate binding domain (SBD), and a 10 kDa C-terminal domain (CTD) which contributes to substrate binding by establishing a lid-like structure over the substrate binding cavity (3–6). Protein refolding by Hsp70s is stimulated by co-chaperones from the Hsp40 and the Hsp110 subfamilies of chaperones (7, 8). Additionally, Hsp70s have a role in clearance of protein aggregates by recruiting members of the Hsp100 family (Hsp104 in yeast) to aggregates, as well as aiding Hsp100 proteins in the disaggregation process (9, 10).

The Stress Seventy sub-family A (SSA) represents the major cytosolic Hsp70 family in yeast and consists of four members: Ssa1-4. Ssa1 and Ssa2 are 97% identical and are both constitutively expressed, while the inducible Ssa3 and Ssa4 are 87% identical to each other and 80% identical to Ssa1/2 (11). The four Ssa isoforms display different spatial/temporal expression patterns but have been thought to be functionally redundant. This notion is supported by the fact that growth can be sustained with the presence of either one of the Ssa isoforms (1). However, there appear to be some functional differences between the isoforms. For example, overproduction of either Ssa3 or Ssa4 as the sole cytosolic Ssa results in increased thermotolerance upon a lethal heat shock, while overproduction of Ssa1 or Ssa2 does not, even though Ssa1 has the most efficient refolding activity of the four isoforms *in vitro* (11, 12). In addition, while Ssa1 modulates prion formation of both *URE3* and *PSI*^+^, Ssa2 does not (11, 13).

Removal of Ssa1 and Ssa2 results in a markedly increased expression of *SSA4* but not *SSA3* (14), yet this strain suffers from increased sensitivity to heat, defects in several protein quality control activities, and accelerated aging (9, 10, 15, 16). Obviously, the enhanced production of Ssa4 cannot fully compensate for the lack of Ssa1 and Ssa2, which suggest that the increased levels of Ssa4 in cells lacking Ssa1/2 are not high enough to fully restore total Ssa levels to those of the wild type cells (as shown in Oling et al., 2014). Another possibility is that Ssa4 cannot carry out some key functions performed by Ssa1 and/or Ssa2. In this work, we approached these possibilities by increasing Ssa4 levels further using a strong constitutive promoter. We found that increased levels of Ssa4 could compensate for the lack of Ssa1 and Ssa2 in almost all PQC processes except the formation of aggregates around the nucleolus and Hsp104-dependent disaggregation, the latter of which was linked to a failure of the nucleotide-binding domain of Ssa4 to recruit Hsp104 to aggregates. The differential role of the Ssa proteins in preventing aggregate formation and aggregate clearance made it possible for us to distinguish which of these processes are most important for longevity assurance.

## RESULTS

### Ssa4 overproduction suppresses heat sensitivity and counteracts formation of protein aggregates in cells lacking both Ssa1 and Ssa2

As previously reported, cells lacking both Ssa1 and Ssa2 display reduced thermotolerance (fig. 1A). By using a strong constitutive promoter (GPD) to drive expression of *SSA4*, we increased the already elevated Ssa4 protein levels in an *ssa1*Δ *ssa2*Δ mutant (16) (fig. S1A) and found that such overproduction of Ssa4 partially suppressed heat sensitivity (fig. 1A). Cells lacking both Ssa1 and Ssa2 accumulate misfolded proteins and protein aggregates already during exponential growth at 30°C, without any applied stress or increased age (16). Using the misfolding proteins guk1-7-GFP and gus1-3-GFP (17), as well as the metacaspase Mca1-GFP, shown to bind and manage protein aggregates (18), as reporters for protein aggregate formation, we could confirm that the *ssa1*Δ *ssa2*Δ mutant accumulated protein aggregates at exponential growth at 30°C to a higher degree than WT cells (fig. 1B, C). Intriguingly, overproduction of Ssa4 suppressed the accumulation of protein aggregates as seen by these three read-outs (fig. 1B, C) and were supported biochemically as Ssa4 overproduction reduced the amounts of insoluble proteins in cells lacking Ssa1/2 (fig. 1D). The suppression of aggregate formation at 30°C was not dependent on the disaggregase Hsp104, as demonstrated with guk1-7-GFP (fig. S1B).

**Figure 1.**
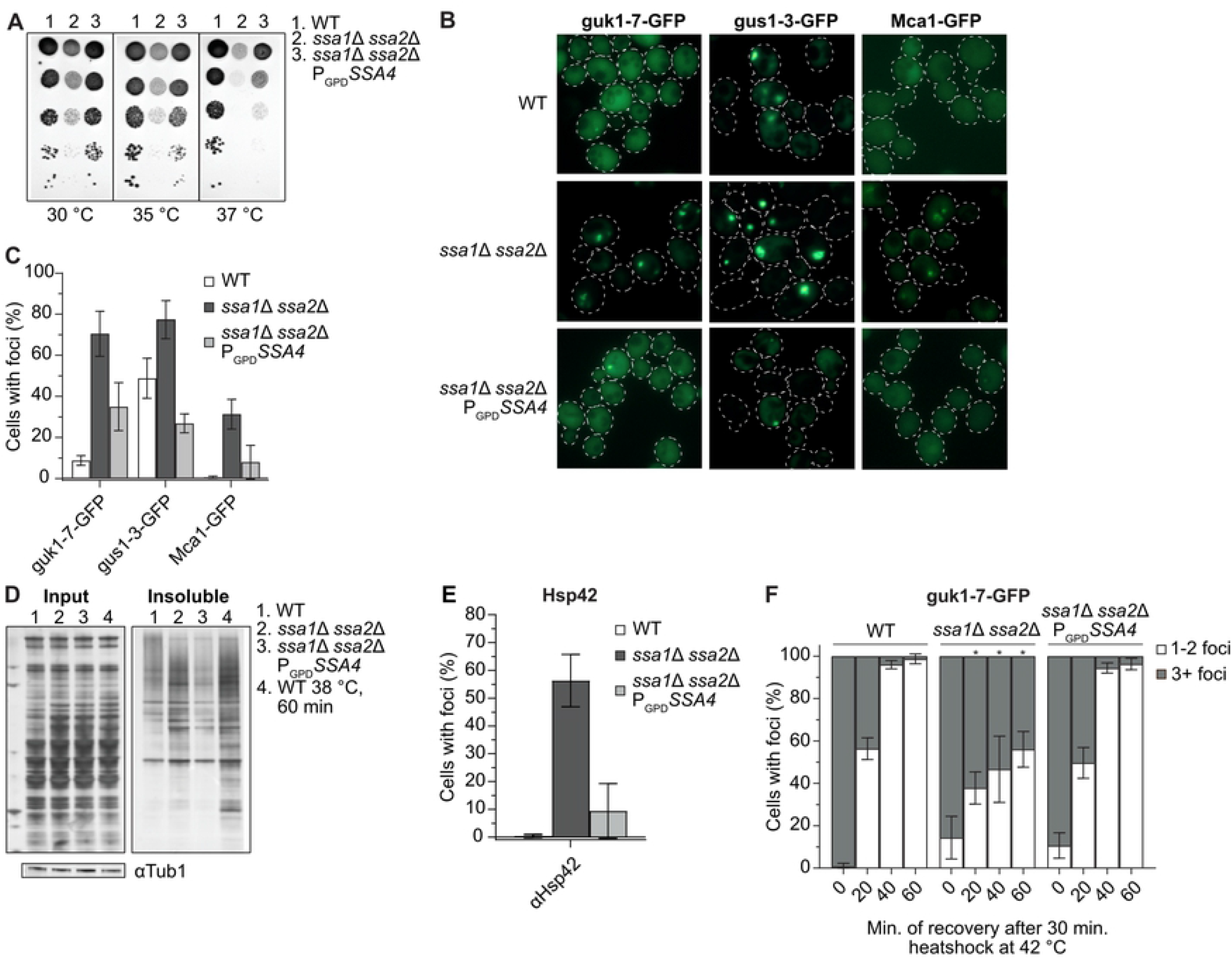
Ssa4 overproduction suppresses heat sensitivity and mitigates protein aggregate accumulation in the absence of Ssa1/2. A) Heat sensitivity test of wild type (WT), and *ssa1*Δ *ssa2*Δ with or without *SSA4* under the control of the GPD-promoter (P_GPD_*SSA4*). The strains were serial diluted 1×10^-1^-1×10^-5^ and incubated on YPD plates for 2 days at the indicated temperatures. B) Fluorescent microscopy images of guk1-7-GFP, gus1-3-GFP and Mca1-GFP in WT, *ssa1*Δ *ssa2*Δ, and *ssa1*Δ *ssa2*Δ P_GPD_*SSA4* background at exponential growth at 30 °C. C) Quantifications of B. Columns represents the mean of 3 biological replicates, *n* = 200. Error bars: +/- S.D. D) Silver stained gel showing total protein input and the insoluble protein fraction of WT, *ssa1*Δ *ssa2*Δ, and *ssa1*Δ *ssa2*Δ P_GPD_*SSA4* during exponential growth at 30 °C (lane 1-3) and heat shocked WT cells (38 °C for 60 minutes, lane 4). Bands below represent immunoblot detection of Tubulin (as a loading control). E) Quantifications of Hsp42 foci in WT, *ssa1*Δ *ssa2*Δ, and *ssa1*Δ *ssa2*Δ P_GPD_*SSA4* using Hsp42 immunofluorescence. Error bars: +/- S.D. F) Guk1-7-GFP foci phenotype of cells with one or more foci after 30 minutes of heat shock at 42 °C and during recovery at 30 °C. Columns represents the mean of 3 biological replicates, *n* = 200. White bars: 1-2 foci per cell with foci, grey bars: 3 or more foci per cell with foci. Note that not all cells have foci in the later time points (see fig. 3D, S1D). Error bars: +/- S.D. * denotes significant difference (*p*<0.05) from the two other strains.

Another marker of protein aggregates is the small heat shock protein Hsp42, which itself is required for sorting proteins into peripheral aggregates (19). Like for guk1-7, gus1-3 and Mca1, we found that Hsp42 resided in aggregates already during exponential growth at 30°C in the *ssa1*Δ *ssa2*Δ mutant, which could be detected through immunofluorescence using antibodies against wild type Hsp42. (fig. 1E, S1C). Overproduction of Ssa4 efficiently reduced Hsp42 foci formation in *ssa1*Δ *ssa2*Δ cells (fig. 1E, S1C). Thus, the data support the notion that aggregate formation, including deposition of Hsp42 into peripheral aggregates, in cells lacking Ssa1/2 can be suppressed by overproducing Ssa4.

Apart from being required for mitigating protein aggregation, Ssa1/2 are also essential for the coalescence of many (3 or more) small aggregates into fewer, larger inclusions upon heat stress (16, 20, 21). We found that Ssa4 overproduction restored the ability of *ssa1*Δ *ssa2*Δ cells to form fewer (1 to 2 foci per cell) inclusions of guk1-7-GFP after heat shock treatment (fig. 1F, S1D, movies S1-3), suggesting that Ssa4 has a role also in the spatial quality control of misfolded proteins in the absence of Ssa1/2.

### Ssa4 overproduction facilitates protein degradation in the absence of Ssa1 and Ssa2

The ability of Ssa4 to mitigate the accumulation of protein aggregates suggest that Ssa4 has a role in the protection against protein aggregate formation and/or the resolution of such aggregates. One strategy for reducing the amount of misfolded proteins in the cytosol is protein degradation by the ubiquitin-proteasome system (UPS) (22). The ΔssCPY*-Leu2-myc (ΔssCL*) construct, which is stabilized in cells with impaired proteasomal degradation and thus confers increased growth on medium lacking leucine, has been used as a read-out for protein degradation by the UPS (16, 23). As previously reported, cells containing the ΔssCL* construct and lacking either Ssa1/2 or the E3 ubiquitin ligase Ubr1 grow better on plates without leucine than wild type cells harboring ΔssCL*. Here, we found that overproduction of Ssa4 restored ΔssCL* degradation in *ssa1*Δ *ssa2*Δ mutant cells, as judged by their poor growth on LEU-plates (fig. 2A). We further confirmed that Ssa4 promotes degradation of misfolded proteins by monitoring the degradation of ΔssCPY*-GFP (ΔssCG*) (24–27) after inhibition of translation by cycloheximide. ΔssCG was stabilized in the *ssa1*Δ *ssa2*Δ mutant and overproduction of Ssa4 destabilized ΔssCG* in the strain lacking Ssa1/2 (fig 2B, C). However, it is unclear if the effect of Ssa4 on protein degradation alone can fully explain the pronounced effect of this chaperone on mitigating aggregate formation (fig. 1B-E).

**Figure 2.**
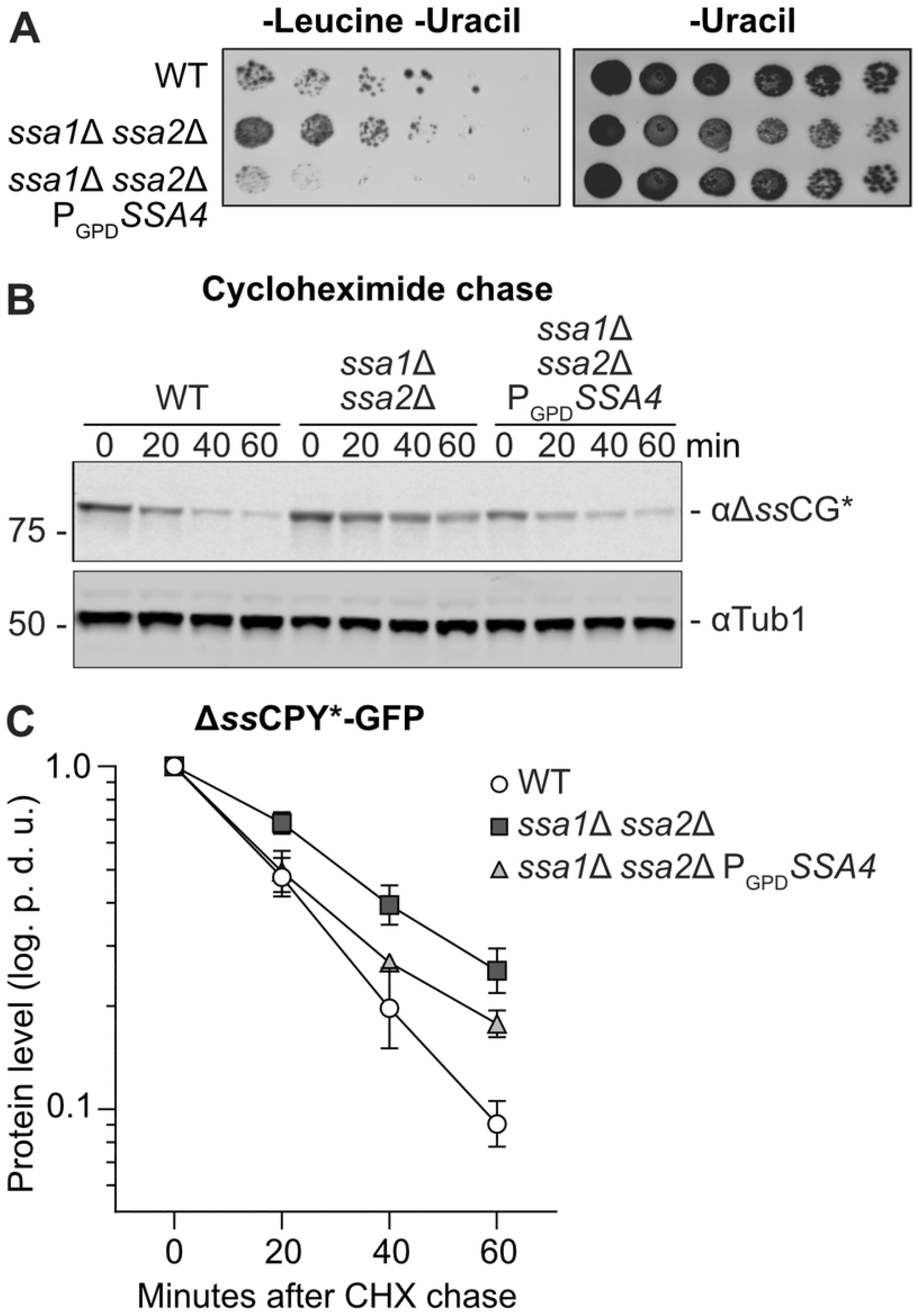
Ssa4 improves degradation of CPY* in the absence of Ssa1/2. A) Growth test of cells expressing ΔssCL* in WT, *ssa1*Δ *ssa2*Δ, and *ssa1*Δ *ssa2*Δ P_GPD_*SSA4* background. The strains were grown on plates lacking leucine and uracil or uracil only (control) for 2-3 days at 30 °C. B) Western blot of Δ*ss*CG* with αGFP antibody after a cycloheximide chase in WT, *ssa1*Δ *ssa2*Δ and *ssa1*Δ *ssa2*Δ P_GPD_*SSA4*. Number panel represents time in minutes after addition of 0.5 mg/ml cycloheximide. C) Quantification of B. Band intensities of the Δ*ss*CG* band were normalized against the corresponding Tub1 band and levels at time point 0 were set to 1 and graphed as the power of 10 logarithm. Error bars represent standard error of the mean of three replicates. p.d.u. = procedure defined unit.

### Ssa4 overproduction does not efficiently promote resolution of heat induced protein aggregates in the absence of Ssa1/2

Resolution of protein aggregates involves the disaggregase Hsp104, which is recruited to protein aggregates in an Ssa1/2-dependent manner (9, 10). Ssa4-GFP localizes to sites of protein aggregation also in the absence of Ssa1/2, and co-localizes with Mca1-RFP, a factor known to localize to proteins aggregates (18), at those sites of aggregation (fig. 3A). Tagging Ssa4 with GFP did not inhibit growth of *ssa1*Δ *ssa2*Δ cells (fig. S1F) demonstrating that the GFP-tag did not disrupt the function of Ssa4 since the loss of *SSA4* in an *ssa1*Δ *ssa2*Δ background is lethal (1). Even though Ssa4 localized to protein aggregates, Ssa4 overproduction did not restore the recruitment of GFP-Hsp104 to protein aggregates in cells lacking Ssa1/2 (fig. 3B, C). Consistently, increased levels of Ssa4 did not fully restore protein aggregate clearance, measured as per cent of Ssa1/2-deficicent cells displaying protein foci of the misfolded guk1-7-GFP after a heat shock (fig. 3D, S1D). This was further confirmed by measuring the GFP intensity ratio between individual foci and the cytosol during recovery (fig. 4A, mov. S1-3) by time-lapse microscopy. This analysis captures the processes in which misfolded proteins are removed from foci, either through disaggregation and refolding and/or degradation. We found that the intensity of guk1-7-GFP foci in the wild type cells was steadily decreasing during recovery in a linear fashion (fig. 3E, left panel, mov. S1). In contrast, there was no such decline in foci intensity in the Ssa1/2 double mutant (fig. 3E, middle panel, mov. S2). Overproducing Ssa4 in the double mutant only partially restored loss of foci intensity in the Ssa1/2 double mutant and the kinetics of the intensity loss was not linear as in wild type cells (fig. 3E, right panel, mov. S3), which hints at a potentially different mechanism underlying the resolution of the foci in these cells (fig 3E, mov. S1-3). Also, as shown in figure S1B, the reduction in aggregate levels achieved by Ssa4 overproduction was not dependent on Hsp104.

**Figure 3.**
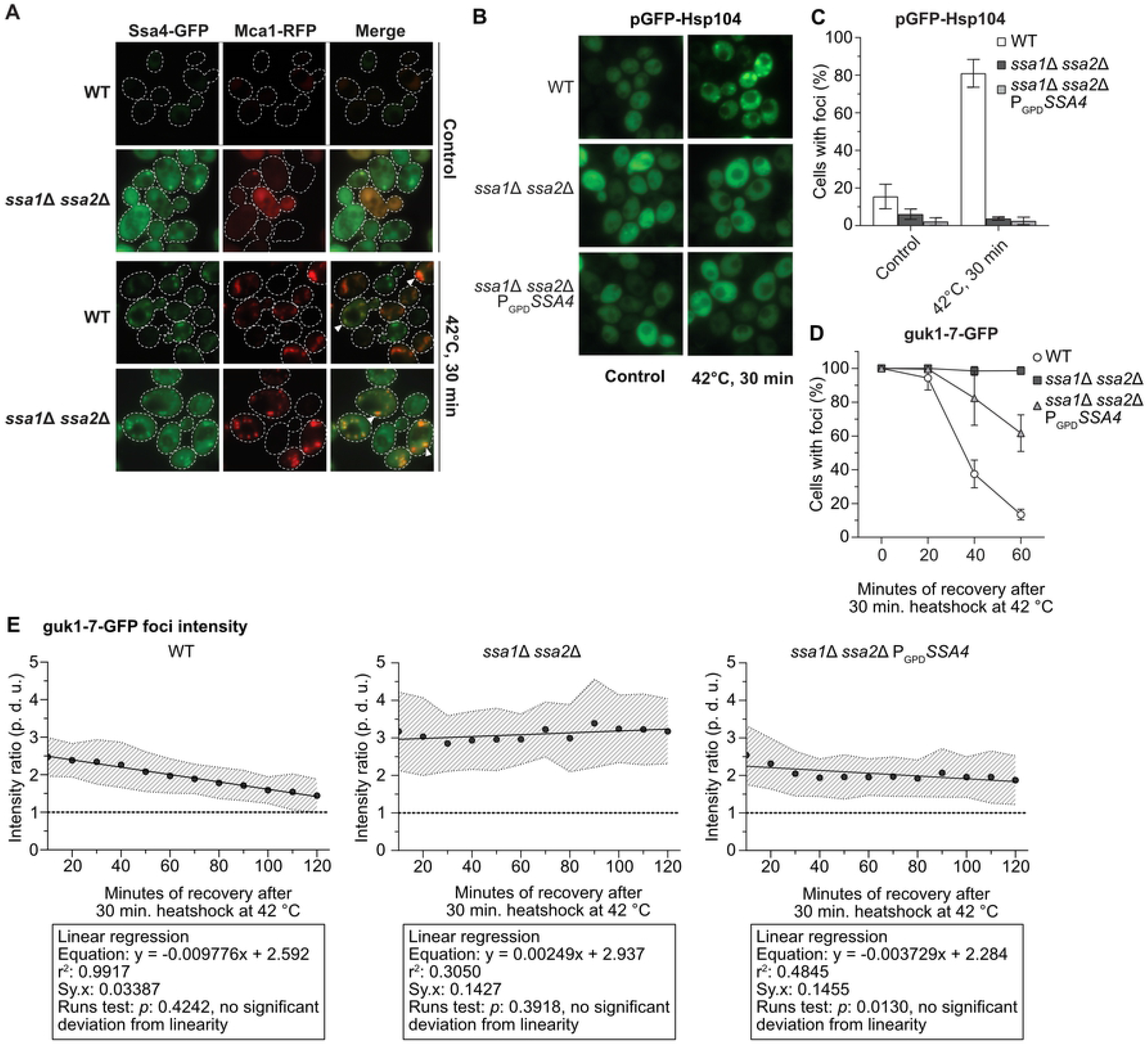
Ssa4 overproduction cannot restore Hsp104 recruitment or protein aggregate resolution in cells lacking Ssa1/Ssa2. A) Fluorescent microscopy images of WT and *ssa1*Δ *ssa2*Δ with Ssa4-GFP and Mca1-RFP under exponential growth at 30 °C and heat shock (42 °C, 30 min) White arrows denotes co-localizing foci. Dashed lines are cell outlines. B) Fluorescent microscopy images of GFP-Hsp104 in WT, *ssa1*Δ *ssa2*Δ, and *ssa1*Δ *ssa2*Δ P_GPD_*SSA4* background under exponential growth at 30 °C and heat shock (42 °C, 30 minutes). C) Quantifications of B. Columns represent the mean of 4 biological replicates, n = 200. White bars: wild type (WT), dark grey bars: *ssa1*Δ *ssa2*Δ, and light grey bars: *ssa1*Δ *ssa2*Δ P_GPD_*SSA4*. Error bars: +/- S.D. D) Quantifications of fluorescent microscopy images of guk1-7-GFP after 30 minutes of heat shock at 42 °C and during recovery at 30 °C. The panel shows the percentage of cells with foci, with the level at time 0 being set to 100 %. Values were normalized to the growth of the culture (measured by the optical density at 600 nm). Symbols represent the mean of 3 biological replicates, n = 200. White circles: wild type (WT), dark grey squares: *ssa1*Δ *ssa2*Δ, and light grey triangles: *ssa1*Δ *ssa2*Δ P_GPD_*SSA4*. Error bars: +/- S.D. E) Ratio between the GFP intensity of foci and cytoplasm in strains with guk1-7-GFP during recovery from 10 to 120 minutes after heat shock (42 °C for 30 minutes). Circles represent the mean ratio of the focus:cytosol intensity at each time point from 20 tracked foci in 20 cells per strain (2 biological replicates), dashed area: +/- S.D. of the mean ratio, dashed lines represents ratio 1:1 = no detectable foci, and the full line is the linear regression of the mean ratios. p. d. u. = procedure defined units. Left panel: WT, Middle panel: *ssa1Δ ssa2Δ*, and Right panel: *ssa1*Δ *ssa2*Δ P_GPD_*SSA4*. The boxes underneath each graph contains the equation of the line from the linear regression, the r^2^ values, standard deviation of the residuals (Sy.x) and the *p*-values of the runs tests performed after the regression.

### Exchanging the nucleotide binding domain (NBD) of Ssa4 allows recruitment of GFP-Hsp104 to protein aggregates

The NBD domain of the *E. coli* Hsp70, DnaK, has been shown to interact with ClpB, a protein of the Hsp100 subfamily and this interaction is important for proper clearance of protein aggregates (28). Similar findings of Hsp70-Hsp100 interactions have been found in *in vitro* studies of yeast-derived Hsp70 and Hsp100 (29, 30), and the IB and IIB ATPase subdomains of the Hsp70 NBD have been pinpointed as the key elements for such interaction between Hsp70 and Hsp104 (30). In order to further elucidate the structure-function relationship between Hsp70 and Hsp104 in aggregate recruitment, we generated chimeric proteins in which each of the three domains (NBD, SBD, or CTD) of Ssa4 was replaced by the Ssa1 counterpart (fig. 4A). This resulted in three different versions of Ssa4, each with one Ssa1-originating domain (NBD_sw_ = NBD swapped Ssa4, SBD_sw_ = SBD swapped Ssa4, CTD_sw_ = CTD swapped Ssa4), that were cloned into centromeric plasmids under the control of the GPD promoter. Introducing either of the three chimeric proteins conferred a robust increase in total Hsp70 protein levels in the *ssa1*Δ *ssa2*Δ background (fig. S2A) and all suppressed the heat sensitivity of the *ssa1*Δ *ssa2*Δ mutant to the same extent as introducing either the wild type *SSA1* or *SSA4* alleles (fig. S2B). We were able to confirm that GFP-Hsp104 recruitment to aggregates was rescued in the *ssa1*Δ *ssa2*Δ background with the plasmid carrying the wild type *SSA1* allele, while the plasmid with *SSA4*, as expected, failed to do so (fig. 4B, C). Interestingly, in cells lacking Ssa1/2, the chimeric version of Ssa4 with the nucleotide binding domain from Ssa1 (pNBD_sw_) was as effective as Ssa1 in the recruitment of GFP-Hsp104 to protein aggregates (fig. 4B, C). No differential effect on the recruitment of GFP-Hsp104 to aggregates was observed when the different alleles were introduced in the WT background (fig. S2C, D). From this data, we conclude that the nucleotide binding domain of Ssa1 is involved in the recruitment of Hsp104 to protein aggregates and that there is a functional difference between the NBDs of Ssa1 and Ssa4. This is in line with *in vitro* data that pinpoint the NBD domain of DnaK as the site of interaction with ClpB in *E. coli* (28).

**Figure 4.**
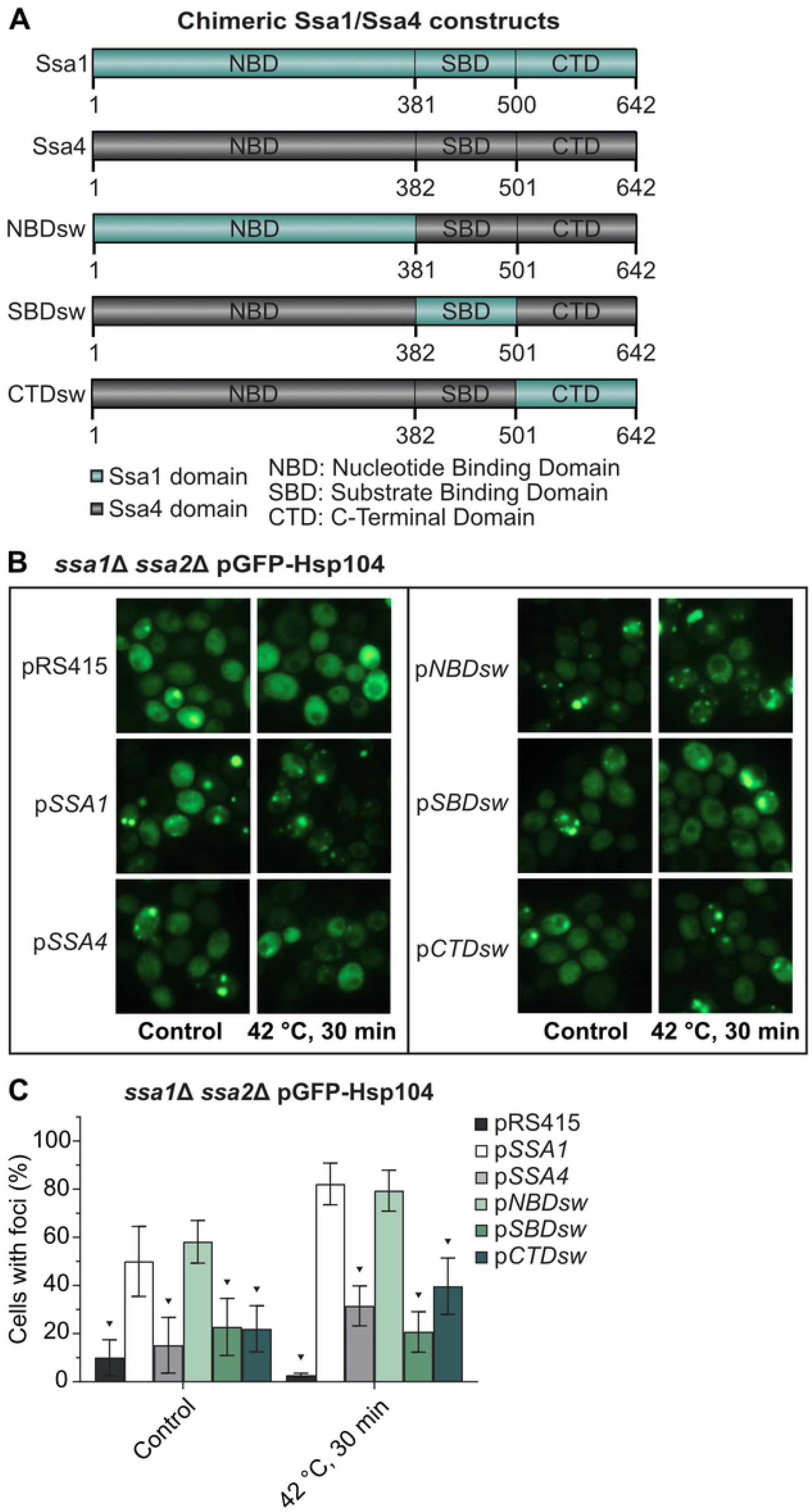
The nucleotide binding domain of Ssa1 but not Ssa4 mediates Hsp104 recruitment to misfolded and aggregated proteins. A) Chimeric Ssa1/Ssa4 constructs used in this study. NBD_sw_ = Nucleotide Binding Domain swapped Ssa4, SBD_sw_ = Substrate Binding Domain swapped Ssa4, CTD_sw_ = C-Terminal Domain swapped Ssa4. B) Fluorescent microscopy images of GFP-Hsp104 in *ssa1*Δ *ssa2*Δ carrying the empty vector (pRS415), wild type Ssa1 and Ssa4 (*pSSA1* and p*SSA4*), and each of the three chimeric versions of Ssa4 (p*NBD*sw, p*SBD*sw and p*CTD*sw) under exponential growth at 30 °C and heat shock (42 °C, 30 min). C) Quantification of cells with foci from B. Columns represent the mean of 6 biological replicates, *n*=193-204. Dark grey: pRS415, white: *pSSA1*, light grey: p*SSA4*, light green: p*NBD_sw_*, green: p*SBD_sw_*, and dark green: p*CTD_sw_*. Error bars: +/- S.D. ▼ symbols above the bars indicate statistically significant difference from p*SSA1* calculated by two-way ANOVA with multiple comparisons between the strains, with Tukey’s post-hoc test, *p* < 0.05. D) Western blot of co-immunoprecipitation with purified Hsp104 protein and cell extracts from *ssa1*Δ *ssa2*Δ *hsp104*Δ cells producing FLAG-tagged versions of either Ssa1 (left), the NBD_sw_ (middle) or the SBD_sw_ (right) chimaeras of Ssa4.

### Formation of ring-like aggregates around the nucleolus is Ssa1/2-dependent and cannot be restored by Ssa4 overproduction

It has been shown previously that the J-protein Sis1 and Ssa2 accumulate in partial ring-like structures around the site of ribosomal RNA transcription and ribosome biogenesis in the nucleolus during heat shock of yeast cells (16, 31), which may be structures akin to those formed due to phase transitions in the nucleolus of mammalian cells subjected to heat stress (32). Using the protein Sik1/Nop56 as a nucleolar reporter (33, 34) and guk1-7 as an aggregate reporter, we found that misfolded proteins also organize into perinucleolar ring structures upon heat shock (fig. 5A, B, mov. S1). The structures persisted through recovery (fig. 5A, C) and eventually reformed into foci that were the last type of guk1-7-GFP aggregates to be resolved after heat shock (mov. S1). Loss of Ssa1/2 inhibited the formation of such structures (fig. 5, mov. S2), while other types of nuclear aggregates were still formed (fig. 5A). The formation of such guk1-7-GFP rings could not be restored by overproducing Ssa4 (fig. 5, mov. S3), and Ssa4-GFP does not organize into such structures upon heat shock (fig. 3A), in contrast to what has been shown previously with Ssa2-GFP (16). Deleting *HSP104* did not abolish the formation of the nucleolar ring-like aggregates, but instead made them more stable during recovery (mov. S4). However, Hsp104 itself does not associate with these structures in the nucleolus (fig. 3B, (31)). Thus, the nucleolar ring-like protein aggregates formed during heat shock are dependent upon Ssa1 and/or Ssa2 to form and Ssa4 cannot compensate for the lack of these chaperones in such aggregate formation.

**Figure 5.**
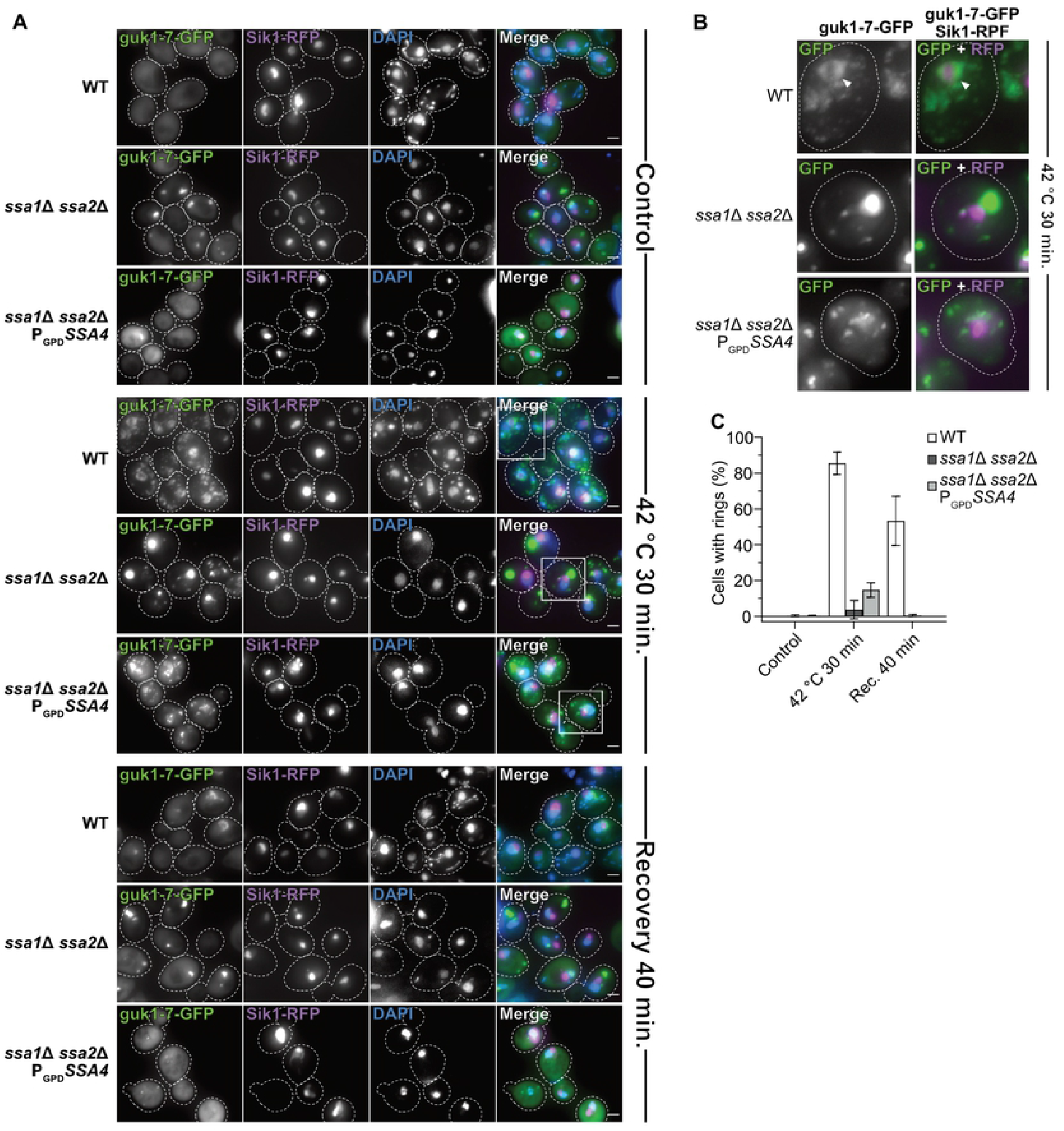
guk1-7-GFP forms an Ssa1/2 ring-like structure around the nucleolus after heat shock. A) Fluorescent microscopy images of guk1-7-GFP, pSik1-RFP and DAPI stain in WT, *ssa1*Δ *ssa2*Δ, and *ssa1*Δ *ssa2*Δ P_GPD_*SSA4* background at control (exponential growth at 30 °C), heat shock (42 °C for 30 minutes) and 40 minutes post-heat shock recovery at 30 °C. Dashed lines represents cellular outlines. White squares shows inserts displayed in B. Green: guk1-7-GFP, magenta: Sik1-RFP, blue: DAPI. B) Zoom in of fluorescent microscopy images of guk1-7-GFP and pSik1-RFP stain in WT, *ssa1*Δ *ssa2*Δ, and *ssa1*Δ *ssa2*Δ P_GPD_*SSA4* background after heat shock (42 °C for 30 minutes). Dashed lines represents cellular outlines. White arrowheads indicates nucleolus-associated ring-like aggregate structure of guk1-7-GFP. Green: guk1-7-GFP, magenta: Sik1-RFP. C) Quantification of percent of cells with guk1-7-GFP ring-like aggregate structures that surrounds the Sik1-RFP signal as shown in A. Columns represent the mean of 3 biological replicates, *n* = 114-201. White: WT, dark grey: *ssa1*Δ *ssa2Δ*, light grey: *ssa1*Δ *ssa2*Δ *P_GPD_SSA4*. Error bars: +/- S.D.

### Ssa4 overproduction fully restores lifespan of short-lived *ssa1*Δ *ssa2*Δ mutant cells

A key question related to PQC and aging is whether it is more important to prevent the formation of aberrant protein conformers, such as aggregates, or to maintain the ability to clear the cells from such aggregates as they form. The differential effects of Ssa4 on aggregate formation and clearance allowed us to approach this question. Specifically, we tested the effect of Ssa4 overproduction on replicative lifespan in cells lacking Ssa1/2 as such overproduction restores folding and degradation of misfolded proteins, mitigates the formation of aggregates, and reinstates inclusion formation, while leaving the cells deficient in the clearance of stress-induced aggregates. Intriguingly, the short lifespan of the *ssa1*Δ *ssa2*Δ mutant was completely restored by Ssa4 overproduction (fig. 6A), suggesting that Hsp104-dependent protein disaggregation is not required to ensure longevity if the degradation and cytosolic spatial quality control mechanisms of the cell are functional.

**Figure 6.**
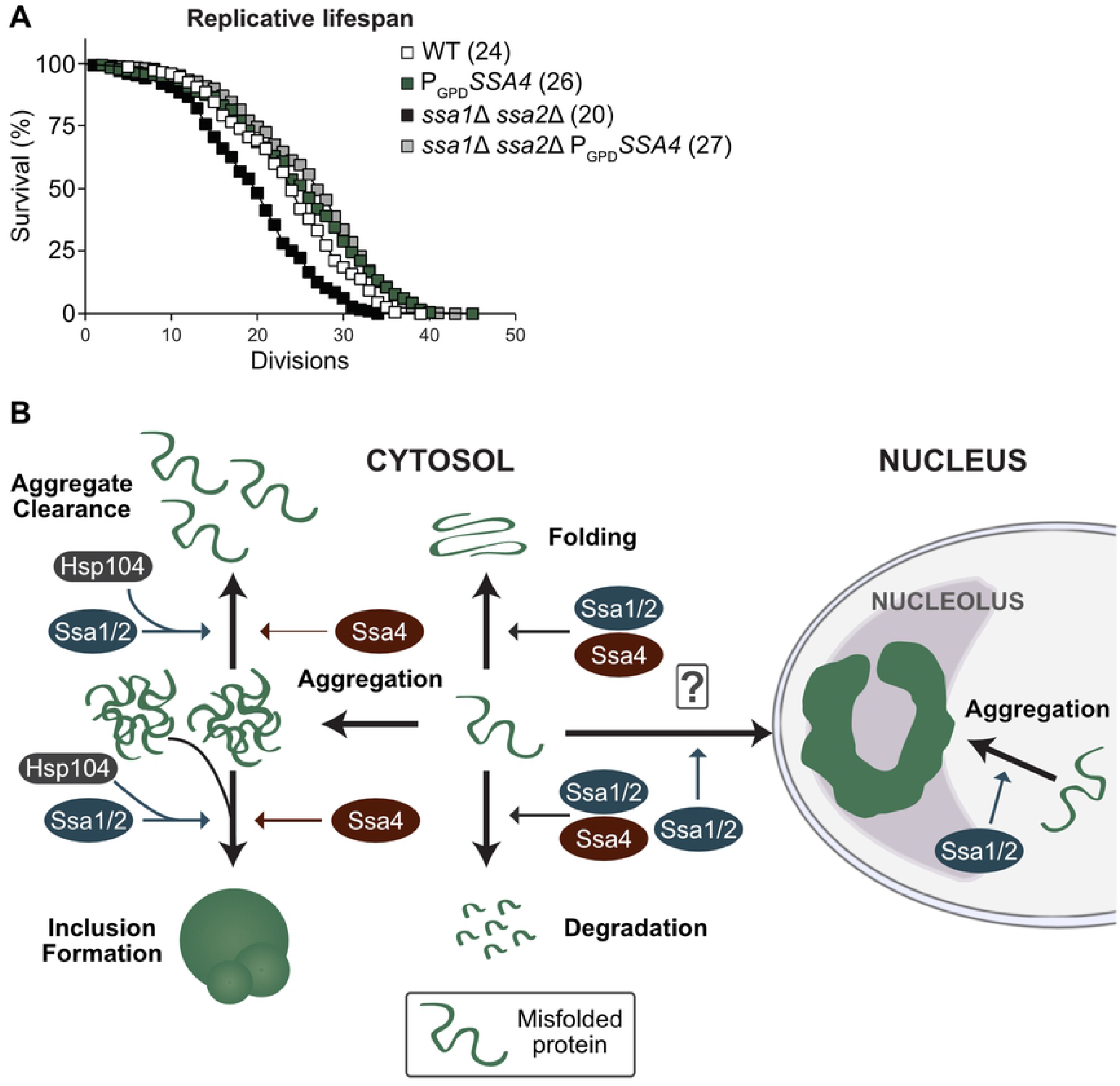
Lifespan analysis and schematic representation of Ssa1 vs. Ssa4 function in PQC. A) Replicative lifespan of WT and *ssa1*Δ *ssa2*Δ strains with or without *SSA4* under the control of the GPD-promoter (P_GPD_*SSA4*). Numbers in parenthesis indicate the mean replicative lifespan of each strain. White squares: WT (*n* = 150), Green squares: P_GPD_*SSA4* (*n* = 179), Black squares: *ssa1*Δ *ssa2*Δ (*n* = 174) and Grey squares: *ssa1*Δ *ssa2*Δ P_GPD_SSA4 (*n* = 321). B) Ssa1/2 (blue) acts on misfolded proteins to promote refolding or degradation. In addition, Ssa1/2 recruits Hsp104 (dark grey) to misfolded, aggregated proteins and aid in the formation of inclusion bodies and nucleolus-associated ring-like aggregates and clearance of protein aggregates. In contrast, Ssa4 (red) can in the absence of Ssa1 and 2 promote refolding and degradation of misfolded proteins as well as aid in the formation of inclusion bodies but does not support the formation of the nucleolus-associated aggregation compartment, nor does it recruit Hsp104 as efficiently to protein aggregates or support efficient clearance of aggregated proteins.

## DISCUSSION

In this paper we have established that ectopically elevating Ssa4 levels in cells lacking the major cytosolic Hsp70 chaperones Ssa1 and Ssa2 can (i) suppress the heat sensitivity (fig. 1A), (ii) counteract protein aggregate accumulation (fig. 1B-E, 6B, S1B), (iii) improve the coalescence of aggregates (fig. 1F, 6B, S1D), (iv) boost degradation of misfolded proteins (fig. 2A-C, 6B) and (v) restore lifespan (fig. 6A). In contrast, Ssa4 overproduction could not efficiently restore Hsp104 recruitment to aggregates and therefore, aggregate clearance remained slow in cells lacking Ssa1/2 (fig. 3B-E, S1E, 6B). This specific inability of Ssa4 to recruit Hsp104 to aggregates appears linked to its nucleotide binding domain, as exchanging this domain to that of Ssa1 fully bestowed Ssa4 with the ability to recruit Hsp104 to misfolded and aggregated proteins (fig. 4B, C). Notably, the previously identified residues in the NBD domain of Ssa1 that have been shown to be important in Ssa1-Hsp104 interactions (L48, Q56, R255, R259, D282, G287 and D289 (30)) are all present in Ssa4 (fig. S2E). If the functional difference between the NBDs of Ssa1 and Ssa4 demonstrated here is due to a difference in the direct interaction between the Ssa in question and Hsp104, or a secondary effect of differential interactions between the NEF partners that compete for the same binding sites (28, 30) remain to be elucidated.

We found also that a nucleolus-associated ring-like structure of the misfolding protein guk1-7-GFP forms upon heat shock and requires Ssa1 and 2 for its formation. Like for aggregate clearance, the formation of this structure cannot be restored by overproduction of Ssa4 in the Ssa1/2 double mutant and the lack of Hsp104 stabilizes the structure after it has formed (mov. S1). A similar structure has been shown before by Öling *et al*. with the use of GFP-tagged Ssa2 in heat shocked cells (16) but, interestingly, we did not see a similar accumulation pattern using Ssa4-GFP in the same conditions (fig 3A). This strengthens the observation that there are functions of Ssa1 or 2 that Ssa4 simply fails to perform. These ring-like structures in the nucleolus of yeast could be a counterpart or analogue to the phase-separated nucleolar compartment for misfolded nuclear proteins described in human embryonic kidney (HEK) 293T cells (32).

A somewhat surprising finding of this work is that Ssa4 overproduction restored inclusion formation in cells lacking Ssa1/2 as Hsp104 is not recruited to aggregates in these cells. Hsp104 has been shown previously to be required for formation of inclusions (20, 35–38) but the data herein indicates that the recruitment of Hsp104 to aggregates may not be a prerequisite for such inclusion formation. Alternatively, inclusion formation in cells lacking Ssa1/2 and overproducing Ssa4 might follow a different route than the one requiring Hsp104 in wild type cells and could possibly lead to differences in spatial locations of inclusions. It should be noted that the role of Hsp104 in longevity assurance is somewhat ambiguous; on the one hand cells lacking Hsp104 age prematurely (39) and lifespan extension by overproducing the metacaspase Mca1 requires Hsp104 (18). On the other hand Hsp104 overproduction on its own does not affect lifespan (40) and lifespan extension by duplicating the gene, *TSA1*, encoding a peroxiredoxin important for Hs70 recruitment to oxidatively damaged proteins, is only marginally affected by deleting *HSP104* (41). In the same vein, there is evidence that aggregation management and protein quality control activities are different during heat shock and ageing (41), and our results here strengthen that idea, since we have shown that not all mechanisms in play during and after heat shock recovery (fig. 3D, 5, S1E) need to be restored in order to maintain a normal life span (fig. 6A), even though they might be necessary for survival at high temperatures (fig. 1A).

While it is clear that Ssa4 is able to compensate for Ssa1/2 in several aspects of PQC and aging, it is also evident that mechanistic differences have evolved within this group of proteins (fig. 6B). Elucidating the individual characteristics of proteins within the same chaperone family may shed light on alternative pathways, such as Hsp104-independent protein inclusion formation and/or protein disaggregation, which will broaden our overall view of different cellular processes related to longevity and proteostasis.

## EXPERIMENTAL PROCEDURES

### Strains and growth conditions

Strains, plasmids, and primers used in this study are listed in Supplementary Tables S1. Strains used in this study are derived from S288C genetic backgrounds. Strains were grown and manipulated by using standard techniques (42). For growth in rich media, strains were grown in yeast extract/peptone/dextrose (YPD) with 2 % glucose. Cells grown in synthetic defined medium were typically grown with yeast nitrogen base plus ammonium sulfate, synthetic complete drop-out mixture and 2% glucose.

### Strain construction

The GPD promoter was amplified from the pYM-N15 plasmid and incorporated upstream of the *SSA4* ORF via homologous recombination in the WT and *ssa1*Δ *ssa2*Δ backgrounds. Deletion mutants were constructed either by PCR-mediated knockout or sporulation. Transformants and dissected spores were verified by PCR. Plasmids with misfolding GPD-tagged proteins (pguk1-7GFP and pgus1-3GFP) were linearized with restriction enzymes (NsiI) and integrated at the *HIS* locus via homologous recombination. The three chimeric *SSA4*-*SSA1* alleles were constructed (GenScript Biotech Corporation, Nanjing, China) and cloned into carrier vectors. The constructs were then amplified with homologous sequences at the 5, and 3’-ends to allow for homologous recombination into the pRS415 vector. The *SSA1* and *SSA4* alleles were amplified from the BY4741 strain with the same homologous sequences. The pRS415 vector was digested with XbaI and HindIII (Thermo Fisher Scientific) and the plasmids were recombined with the constructs in wild type yeast and then extracted and transformed into *E. coli* to allow for plasmid production. The final vectors were digested with HindIII and SacI (Thermo Fisher Scientific) for insert size control and then sequenced for verification prior to transformation into the intended host strains.

### Heat-sensitivity assay

Cultures were grown to mid-exponential phase at 30°C, and serially diluted (tenfold) to an optical density (38) of A_600_ = 10^-1^–10^-5^. Dilutions were spotted onto YPD or synthetic defined medium plates and incubated at the indicated temperatures for 2-3 days prior to imaging.

### Replicative lifespan analysis

Lifespan analyses were performed as previously described (Kaeberlein et al, 1999, Erjavec et al, 2007). Briefly, cells were grown overnight in YPD, diluted, plated, and allowed to recover on YPD plates before being assayed for RLS which was performed using a micromanipulator (Singer instruments) to remove daughters from mother cells. Replicative lifespan analysis was performed independently at least three times for each strain. Statistical analysis was performed using unpaired Mann-Whitney U-test in GraphPad Prism^®^ 7.01.

### Stress-treatments used to induce protein aggregation

Cells were grown to mid-exponential phase at 30°C and then shifted to the indicated temperatures. For visualization of protein aggregate formation, cells were kept at indicated temperatures and samples were taken over time, and cells were scored as either having no or one or more protein aggregate foci. For resolution of protein aggregates and aggregate coalescence after heat shock, cells were shifted back to 30°C after heat shock and samples were taken over time. For the resolution of protein aggregate foci cells were scored as either having none or one or more protein aggregate foci. The OD of each culture was measured at each sampling time point for normalization. Protein aggregate coalescence/inclusion formation during recovery was scored in three categories: foci-free (0 foci), class I/II (1-2 foci) and class III (3 or more foci). All cells in the foci-free category was omitted and the remaining cells with at least one foci were used in the analysis. Images were analyzed with ImageJ (Fiji package) (43). Statistical analyses were performed in GraphPad Prism^®^ 7.01 (GraphPad Software LLC) and Microsoft^®^ Excel^®^ 2016 (Microsoft Corporation).

### GFP-Hsp104 expression from *GAL1*-promoter

Cultures were grown over the day in synthetic defined medium lacking histidine (CSM-HIS) or leucine and histidine (CSM-LEU-HIS) with 2% galactose. Cultures were resuspended in fresh medium with 2% galactose and grown to mid-exponential phase over the night. Sample of mid-exponential phase cells were taken before and after heat shock (42 °C, 30 minutes).

### DAPI-stain of live cells

Cells were cultured, treated and sampled as described above. Prior to imaging, 10 μg/ml of 4’,6-diamidino-2-phenylindole (DAPI) in MilliQ H_2_O was added to each cell sample and incubated in darkness for 5 minutes before pelleting by centrifugation and subsequent imaging.

### Fluorescence microscopy

Images were obtained using a conventional fluorescence microscope, a Zeiss Axio Observer.Z1 inverted microscope with Apotome and Axiocam 506 camera, and a Plan-Apochromat 100x/1.40 Oil DIC M27 objective.

### Isolation of insoluble protein

Insoluble protein aggregates were isolated as described (Rand and Grant, 2006), with minor adjustments. The protocol was scaled up to suit the harvesting of 35 A600 units of cells. Pefabloc SC (Roche) was used instead of phenylmethylsulfonyl fluoride in the lysis buffer. After freezing and thawing the cells they were immediately disrupted by a minibead beater (Fastprep) at 5.5 m/s for 4×30 sec at 4°C. Cells were kept on ice for 5 min between each beating. Removal of intact cells was accomplished by centrifugation at 1180 x g for 15 min. Protein concentration was determined using Pierce 660 nm Assay Reagent (Thermo Scientific) and adjusted before isolation of protein aggregates as well as verified by running reduced samples on Criterion XT Precast Gel 4-12% Bis-Tris (BioRad) and staining with Coomassie Brilliant Blue R-250 staining solution (BioRad). The subsequent centrifugations were all done at 21130 x g for 20 min. Insoluble fractions were resuspended in detergent washes by 5 x 5 sec sonication at amp 40 with 15 sec rest between intervals using Q700 Sonicator (QSonica, LLC. Newtown, USA). The insoluble fractions were added to reduced protein loading buffer, loaded on Criterion XT Precast Gel 4-12% Bis-Tris (BioRad) and visualized by silver staining with Pierce^®^ Silver Stain Kit (Thermo Scientific).

### Growth test for cells expressing ΔssCL*

Overnight pre-cultured cells were diluted to an O.D. of A_600_=0.5, spotted in a fivefold dilution series on CSM-URA or CSM-URA-LEU plates and incubated at 30°C for 3–4 days.

### Protein extraction, gel electrophoresis and western blot

Cells were cultured overnight and then diluted down to an O.D. of A_600_= 0.1 and allowed to grown to an O.D. of A_600_=0.5 before harvest. For denaturing gel electrophoresis, proteins were extracted as described by (44) with a few modifications. Briefly, 1 O.D. unit of cells were lysed in 200 μl of 0.2 M NaOH for 20 minutes on ice followed by centrifugation for 1 minute at 13000 rpm, 4 °C. The protein pellet was resuspended in 50 μl per O.D. unit of cells in a 1X Laemmli/8 M Urea/ 2.5 % β-mercaptoethanol sample buffer and incubated for 10 minutes at 70 °C prior to SDS-PAGE using 4-12 % Bis-Tris gels (Invitrogen NuPAGE^®^, Thermo Scientific) in a MOPS buffer (Invitrogen NuPAGE^®^, Thermo Scientific) according to the manufacturers instruction. The gels were blotted onto an Immobilion-P PVDF membrane, 0.45 μm pore size, (MERCK Millipore) in a Tris-glycine-methanol transfer buffer overnight and detected with the following antibodies: 1:5000 monoclonal mouse anti-Hsp70 [C92F3A-5] (Abcam Cat# ab47455, RRID:AB_881520) (abcam), 1:15000 monoclonal mouse anti-Pgk1 (Thermo Fisher Scientific Cat# 459250, RRID:AB_2532235). The primary antibodies were detected using 1:20000 goat anti-mouse or rat IgG (H+L) IRDye^®^ 680LT or 800CW (mouse 680LT: LI-COR Biosciences Cat# 926-68020, RRID:AB_10706161, mouse 800CW: LI-COR Biosciences Cat# 926-32210, RRID:AB_621842, rat 680LT: LI-COR Biosciences Cat# 926-68029, RRID:AB_10715073, rat 800CW: LI-COR Biosciences Cat# 926-32219, RRID:AB_1850025), the blots were scanned using the Odyssey^®^ Near Infra-red imaging system (LI-COR) and the images were analyzed using ImageJ.

### Cycloheximide chase

Logarithmically growing cells expressing ΔssCG* were harvested and resuspended in complex media. Protein translation was shut off by addition of cycloheximide (PubChem CID: 6197) (0.5 mg/ml final concentration). 2 O.D. units of cells were taken at indicated time-points, protein samples were prepared with minor changes as described above. 8 M Urea loading dye was supplemented with complete protease inhibitor (Roche). 100 μl Urea loading dye were used to resuspend cell pellet after NaOH treatment. 15 μl of the protein extracts were loaded on a 4-12% gradient 26 well Criterion XT Bis-Tris Protein gel (Bio-Rad).

Gels were transferred on PVDF membrane. Membranes were blocked with Odyssey blocking buffer (Li-COR) followed by incubation with antibodies specific for GFP (mouse monoclonal anti-GFP [7.1 and 13.1] (Sigma-Aldrich Cat# 11814460001, RRID:AB_390913)) and Tub1 (rat monoclonal anti-tubulin alpha, (Bio-Rad Cat# MCA77G, RRID:AB_325003)) (1:10000 dilution each in PBS-T). The primary antibodies were detected and the membranes scanned as described above.

### Immunofluorescence

Cells were fixed in 9 parts 0.1 mM KPO_4_, 500 mM MgCl_2_ and 1 part 37 % formaldehyde for 1h at 30 °C, washed once in 0.1 M KPO_4_, 500 μM MgCl_2_ and once in 0.1 M KPO_4_, 500 μM MgCl_2_, 0.1 M Sorbitol (sorbitol solution). Spheroplasts were prepared by incubating approximate 3 O.D. units of fixed cells at 35 °C with 0.6 mg/ml Zymolase in sorbitol solution. Spheroplasts were gently washed in sorbitol solution before being spotted on polylysin-coated slides and permeabilized swiftly with 1 % Triton X-100 in BSA-PBS (1 % BSA). Blocking was done in BSA-PBS. To avoid unspecific binding of primary antibody, polyclonal rabbit anti-Hsp42, 1:2000 (gift from Prof. Johannes Buchner, Germany) in PBS-BSA, was pre-incubated on a western blot membrane with transferred BY4741 *hsp42*Δ::*kanMx* lysate for 1.5 h at room temperature (RT) before antibody solution was added to slides. Slides were incubated with 1:500 goat anti-Rabbit Alexa488 (Thermo Fisher Scientific Cat# A-11034, RRID:AB_2576217), washed, incubated with 1 μg/ml DAPI for 10 min at RT and washed before mounting media was added and slides were sealed with nail polish. Mounting media was prepared by adding 100 mg p-phehylenediamine in 10 ml PBS, pH was adjusted to 8.0 with 0.5M Sodium carbonate buffer (pH 9.0) and volume was brought to 100 ml by addition of glycerol.

### Time lapse microscopy

Cells were grown in YPD medium to an O.D._600_ between 0.3 and 0.5 at 30 °C after which they were heat shocked at 42 °C for 30 minutes. The heat shocked samples were washed with PBS and then placed onto agar pads with complete synthetic media supplemented with YNB and 2 % glucose mounted on microscope slides. The samples were imaged at 30 °C on a Zeiss Axio Observer.Z1 inverted microscope with Apotome and Axiocam 506 camera, and a Plan-Apochromat 100x/1.40 Oil DIC M27 objective, with the following incubation equipment to maintain temperature: TempModule S1 (Zeiss), Y module S1 (Zeiss), Temperable insert S1 (Zeiss), Temperable objective ring S1 (Zeiss), and Incubator S1 230 V (Zeiss). 10 *Z*-stack images were captured at four distinct coordinates every 10 minutes for 120 minutes. The time lapse captures were processed in ImageJ (Fiji package) (43) using the Bleach Correction with Simple Ratio (45) and the Linear Stack Alignment with SIFT (46) plugins. The analysis was performed in ImageJ by measuring the Integrated Density (IntDen_ROI_) and Area (A_ROI_) of a focus and a same-sized spot of the cytoplasm for each focus and time point. Mean fluorescence measurements (M_B_) were also taken of the cell-free background. The corrected cellular compartment fluorescence for each focus and associated foci-free cytosol was then calculated for each focus and time point with the following equation: IntDen_ROI_ – (A_ROI_ * M_B_). After that, the ratio between the individual focus and corresponding cytosol was calculated for each time point. The linear regression and statistical analysis were carried out using GraphPad Prism^®^ 7.01 (GraphPad Software LLC).

## ACKNOWLEDGEMENTS

The authors would like to thank Johannes Buchner for the generous gift of the rabbit anti-Hsp42 antibody, Charles Boone for strains and Per Widlund for the gus1-3-GFP and guk1-7-GFP plasmids used in this study.

## SUPPLEMENTAL MOVIE CAPTIONS

**Movie S1.** Time lapse microscopy of BY4741 guk1-7-GFP cells during recovery at 30 °C after heatshock at 42 °C for 30 minutes. The movies show 12 images captured every 10 minutes from 10 to 120 minutes after the heat shock was completed.

**Movie S2.** Time lapse microscopy of *ssa1*Δ *ssa2*Δ guk1-7-GFP cells during recovery at 30 °C after heatshock at 42 °C for 30 minutes. The movies show 12 images captured every 10 minutes from 10 to 120 minutes after the heat shock was completed.

**Movie S3.** Time lapse microscopy of *ssa1*Δ *ssa2*Δ P_GPD_*SSA4* guk1-7-GFP cells during recovery at 30 °C after heatshock at 42 °C for 30 minutes. The movies show 12 images captured every 10 minutes from 10 to 120 minutes after the heat shock was completed.

**Movie S4.** Time lapse microscopy of *hsp104*Δ guk1-7-GFP cells during recovery at 30 °C after heatshock at 42 °C for 30 minutes. The movies show 12 images captured every 10 minutes from 10 to 120 minutes after the heat shock was completed.

## SUPPLEMENTAL FIGURE CAPTIONS

**Figure S1.** A) Left: Cut-outs of silver stained 2-dimensinal polyacrylamide gels showing Ssa1, Ssa2 and Ssa4 in protein extracts from WT, P_GPD_*SSA4*, *ssa1*Δ *ssa2*Δ and *ssa1*Δ *ssa2*Δ P_GPD_*SSA4* cells. Black arrows indicate the named proteins. Right: Quantification the gels shown to the left with respect to Ssa4 levels relative to Act1. WT is set to 1. B) Quantification of number of foci per cell of guk1-7-GFP in WT, *ssa1*Δ *ssa2*Δ and *ssa1*Δ *ssa2*Δ P_GPD_*SSA4* with (left) or without (right) *HSP104*. C) Fluorescent microscopy images of Hsp42 in WT, *ssa1*Δ *ssa2*Δ, and *ssa1*Δ *ssa2*Δ P_GPD_*SSA4* by anti-Hsp42 antibody detection D) Fluorescent microscopy images of guk1-7-GFP cells after heat shock (42 °C for 30 minutes) and during recovery at 30 °C. Shown are images representative of three biological replicates. Images were generated as a maximum projection of one representative slice from a *Z*-stack using ImageJ 1.50i (Rasband, 1997) with 64-bit Java 1.8.0_77. E) Heat sensitivity test of WT and *ssa1*Δ *ssa2*Δ with or without GFP-tagged Ssa4.

**Figure S2.** A) Representative western blot of SDS-PAGE gel with anti-Hsp70 antibody and anti-Pgk1 antibody as loading control (left). Quantification of immunoblotted SDS-PAGE gel with respect to total Hsp70 (right). Values have been normalized to Pgk1 as loading control and the level in WT at steady state set to 1. Bars represent one representative replicate out of two. White with black dots: WT pRS415, dark grey: *ssa1*Δ *ssa2*Δ pRS415, white: *ssa1*Δ *ssa2*Δ pSSA1, light grey: *ssa1*Δ *ssa2*Δ pSSA4, light green: *ssa1*Δ *ssa2*Δ pNBD_sw_, green: *ssa1*Δ *ssa2*Δ pSBD_sw_, and dark green: *ssa1*Δ *ssa2*Δ pCTD_sw_. B) Heat sensitivity test of wildtype (WT), and *ssa1*Δ *ssa2*Δ with the empty vector (pRS415), wild type Ssa1 and Ssa4 (*pSSA1* and p*SSA4*), and each of the three chimeric versions of Ssa4 (p*NBD_sw_*, p*SBD_sw_* and *pCTD_sw_*). The strains were serial diluted 1×10^-1^-1×10^-5^ and plated on synthetic medium agar plates for 3 days at the indicated temperatures. C) Fluorescent microscopy images of GFP-Hsp104 in wild type cells carrying the empty vector (pRS415), wild type Ssa1 and Ssa4 (*pSSA1* and p*SSA4*), and each of the three chimeric versions of Ssa4 (*pNBD_sw_, pSBD_sw_* and *pCTD_sw_*), under exponential growth at 30 °C (control) and heat shock (42°C, 30 minutes). D) Quantification of cells with foci from B. Columns represent the mean of 3 biological replicates, *n* = 200-209. Dark grey: pRS415, white: pSSA1, light grey: pSSA4, light green: pNBD_sw_, green: pSBD_sw_, and dark green: pCTD_sw_. Error bars: +/- S.D. E) Global alignment of Ssa1 and Ssa4. Conservation bar indicates the number of conserved physiochemical properties between the residues where * = conserved residue, + = all properties conserved, 9 - 1 = almost all to one property conserved, - = insertion/deletion/gap. Arrows (↓) mark residues previously identified as important in Ssa1-Hsp104 interactions (30). Shaded boxes over residue codes indicate domain demarcations; light grey: NBD, grey: SBD, dark grey: CTD

## Notes

### Competing Interest Statement

D. Ö. is employed by AstraZeneca Mölndal, Sweden
F.E. is employed by AstraZeneca Mölndal, Sweden
S.H. is employed by Cochlear Nordic AB, Mölnlycke, Sweden

### Summary of Updates

Updated with supplemental data and author ORCID id's.

