## Supplementary figures and images for "Differential role of cytosolic Hsp70s in longevity assurance and protein quality control"

### Supplemental figure 1

#### SUPPLEMENTAL FIGURE 1

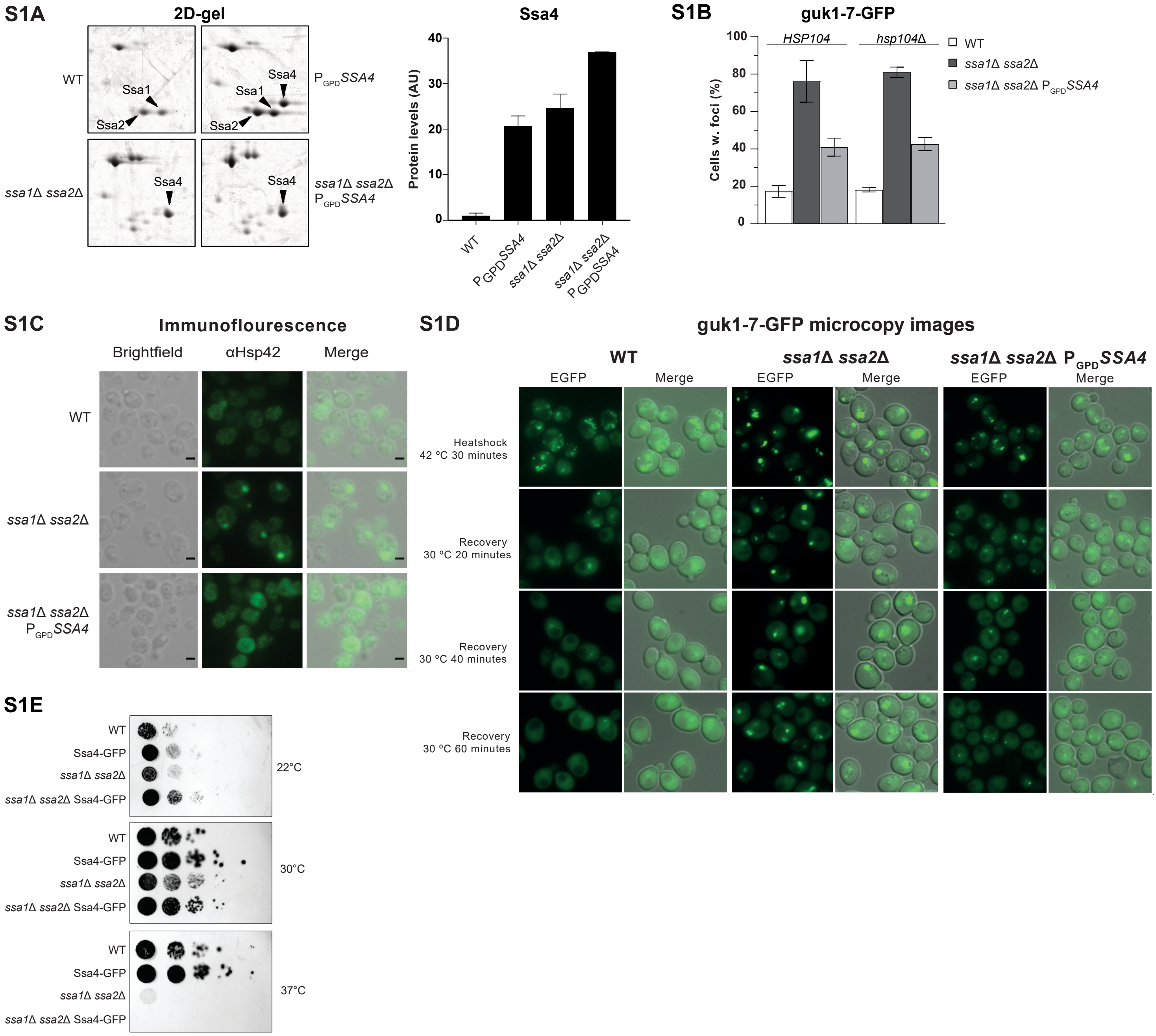


SUPPLEMENTAL FIGURE 1

### Supplemental figure 2

##### SUPPLEMENTAL FIGURE 2

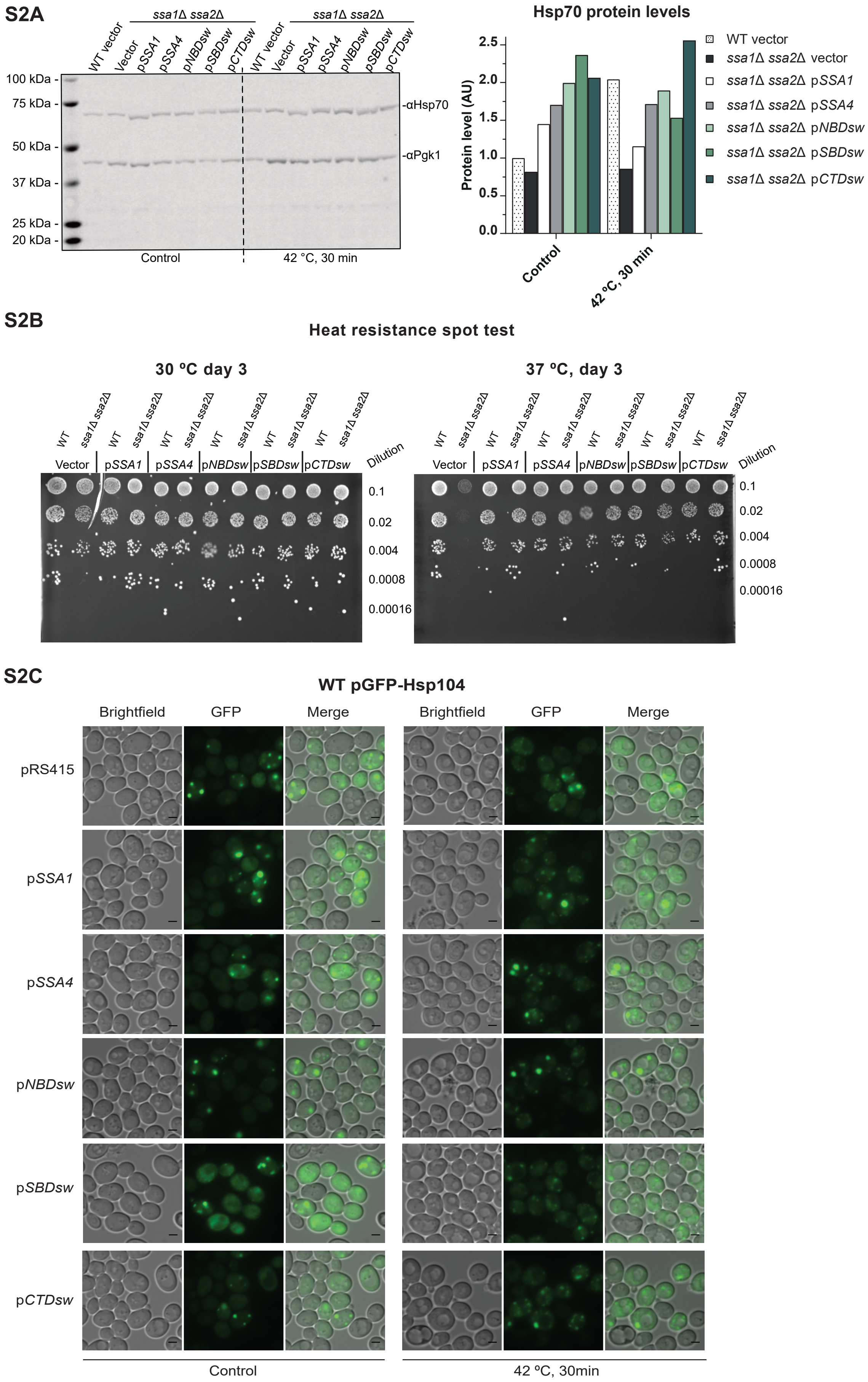

#### SUPPLEMENTAL FIGURE 2

## S2D

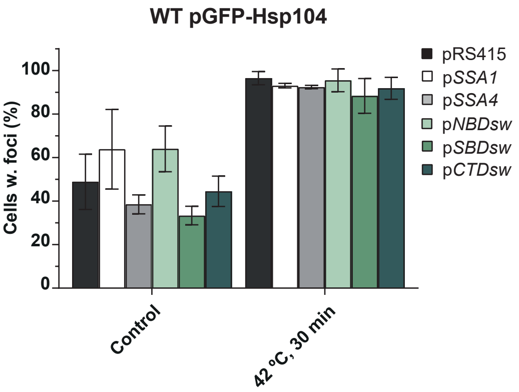


##### S2E Global alignment of Ssa1 and Ssa4

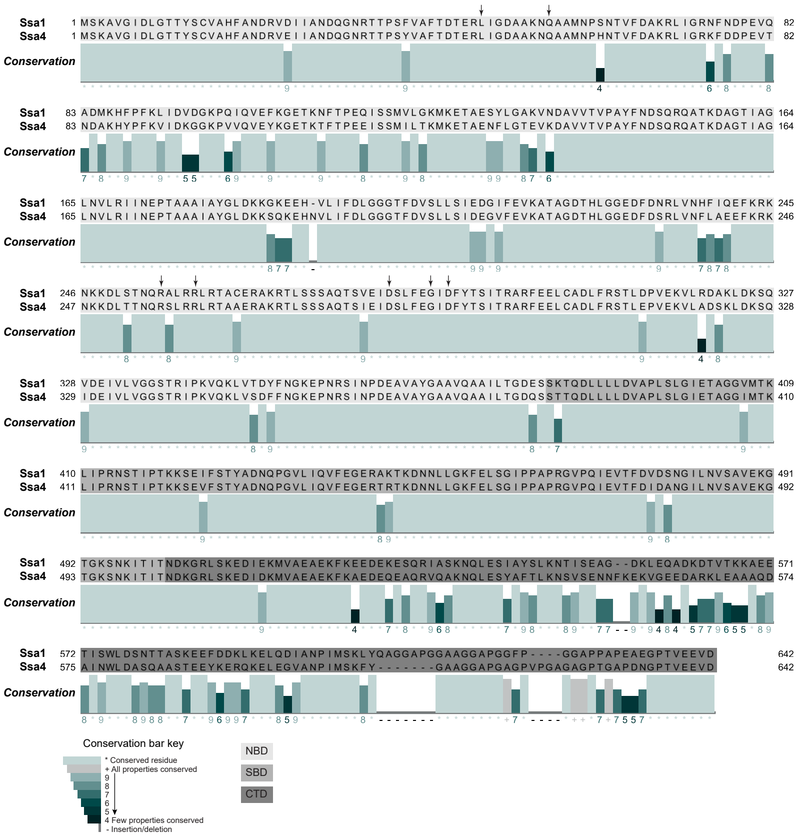
