## Supplemental materials and methods for "Differential role of cytosolic Hsp70s in longevity assurance and protein quality control"

### SUPPLEMENTAL MATERIAL

#### SUPPLEMENTAL MATERIAL AND METHODS

**Two-dimensional polyacrylamide gel electrophoresis (2D-PAGE).** Cell harvesting, protein extraction, and 2D-PAGE were performed as described previously (Maillet et al., 1996, Blomberg, 2002). For the first dimension, strips ranging from pH 3–5.6 were used (GE Healthcare). The second-dimension gels contained 7 % polyacrylamide. Total protein was visualized using silver staining. Staining was performed accordingly: fixation in 50% ethanol (v/v) plus 10% acetic acid for 2 h followed by washing in MilliQ water 3 × 20 min. Gels were sensitized in 500 ml DTT solution (5.2 mg/l) for 30 min. Subsequently, gels were incubated for 30 min in 500 ml silver nitrate solution (2 g/l). Gels were washed in 500 ml MilliQ water for 30 s, water poured off and 200 ml sodium bicarbonate (34.7 g/l) with formaldehyde (0.5 g/l) was added until a color development was observed. The solution was discarded and 500 ml fresh sodium bicarbonate with formaldehyde solution was added. Development of gels was stopped after approximately 5–6 min. Silver-stained gels were scanned, aligned, and quantified with ImageJ. Ssa1–4 proteins were identified by visual matching with existing yeast 2-D maps.

**Protein alignment.** Protein alignment of Ssa1 (SGD, 2018a) and Ssa4 (SGD, 2018b) were performed on amino acid sequences obtained from the S288C reference strain. Protein alignments were performed in Geneious 8.0.5 ® (Biomatters Ltd, New Zealand) by a global alignment using the Needleman-Wunsch algorithm (Needleman and Wunsch, 1970) with BLOSUM62 cost matrix (Henikoff and Henikoff, 1992), and gap open penalty: 10, gap extension penalty: 0.5. Image was generated using Jalview 2 (Waterhouse et al., 2009), conservation scores calculated as described by Livingstone and Barton (Livingstone and Barton, 1993).

**TABLE S1: Strain list**

| Figures | Name | Genotype | Reference |
|---|---|---|---|
| 1A, D, E<br>2<br>3B, C<br>6A<br>S1A, C, E<br>S2A-D | BY4741/WT | Mat **a** *his3Δ1 leu2Δ0 met15Δ0 ura3Δ0* | (Brachmann et al., 1998) |
| 6A<br>S1A | $P_{GPD}$*SSA4* | Mat **a** *his3Δ1 leu2Δ0 met15Δ0 ura3Δ0*<br>*natMX6:*$P_{GPD}$*-SSA4* | This study |
| 1A, D, E<br>2<br>3B, C<br>4B, C<br>6A<br>S1A, C, E<br>S2A, B | *ssa1Δ ssa2Δ* | Mat **a** *his3Δ1 leu2Δ0 met15Δ0 ura3Δ0*<br>*ssa1Δ::hph ssa2Δ::kanMX4* | (Oling et al., 2014) |
| 1A, D, E<br>2<br>3B, C<br>6A<br>S1A, C | *ssa1Δ ssa2Δ*<br>$P_{GPD}$*SSA4* | Mat **a** *his3Δ1 leu2Δ0 met15Δ0 ura3Δ0*<br>*ssa1Δ::hph ssa2Δ::kanMX4*<br>*natMX6:*$P_{GPD}$*-SSA4* | This study |
| 1B, C, F<br>3D, E<br>5<br>S1B, D<br>M1A | WT guk1-7-GFP | Mat **a** *his3*::pRS403-*guk1-7-EGFP::HIS3*<br>*leu2Δ0 met15Δ0 ura3Δ* | This study |
| 1B, C, F<br>3D, E<br>5<br>S1B, D<br>M1B | *ssa1Δ ssa2Δ* guk1-7-GFP | Mat **a** *his3*::pRS403-*guk1-7-EGFP::HIS3*<br>*leu2Δ0 met15Δ0 ura3Δ*<br>*ssa1Δ::hph ssa2Δ::kanMX4* | This study |
| 1B, C, F<br>3D, E<br>5<br>S1B, D<br>M1C | *ssa1Δ ssa2Δ*<br>$P_{GPD}$*SSA4* guk1-7-GFP | Mat **a** *his3*::pRS403-*guk1-7-EGFP::HIS3*<br>*leu2Δ0 met15Δ0 ura3Δ*<br>*ssa1Δ::hph ssa2Δ::kanMX4*<br>*natMX6:*$P_{GPD}$*-SSA4* | This study |
| 1B, C | WT gus1-3-GFP | Mat **a** *his3*::pRS403-*gus1-3-EGFP::HIS3* | This study |

| | | | |
|---|---|---|---|
| | | *leu2Δ0 met15Δ0 ura3Δ* | |
| 1B, C | *ssa1Δ ssa2Δ gus1-3-* GFP | Mat **a** *his3*::pRS403-*gus1-3-EGFP::HIS3 leu2Δ0 met15Δ0 ura3Δ ssa1Δ::hph ssa2Δ::kanMX4* | This study |
| 1B, C | *ssa1Δ ssa2Δ* $P_{GPD}$*SSA4* gus1-3-GFP | Mat **a** *his*::pRS403-*gus1-3-GFP::HIS3 leu2Δ0 met15Δ0 ura3Δ ssa1Δ::hph ssa2Δ::kanMX4 natMX6:*$P_{GPD}$*-SSA4* | This study |
| 1B, C | Mca1-GFP | Mat **a** *his3Δ1 leu2Δ0 met15Δ0 ura3Δ0 MCA1-EGFP::HIS6* | (Huh et al., 2003) |
| 1B, C | *ssa1Δ ssa2Δ* Mca1-GFP | Mat **a** *his3Δ1 leu2Δ0 met15Δ0 ura3Δ0 MCA1-EGFP::HIS6 ssa1Δ::hph ssa2Δ::kanMX4* | (Hill et al., 2014) |
| 1B, C | *ssa1Δ ssa2Δ* $P_{GPD}$*SSA4* Mca1-GFP | Mat **a** *his3Δ1 leu2Δ0 met15Δ0 ura3Δ0 MCA1-EGFP::HIS6 ssa1Δ::hph ssa2Δ::kanMX4 natMX6:*$P_{GPD}$*-SSA4* | This study |
| 3A | Ssa4-GFP Mca1-RFP | Mat **a** *his3Δ1 leu2Δ0 met15Δ0 ura3Δ0 SSA4-GFP::HIS3 MCA1-RFP::LEU2* | This study |
| 3A | *ssa1Δ ssa2Δ* Ssa4-GFP Mca1-RFP | Mat **a** *his3Δ1 leu2Δ0 met15Δ0 ura3Δ0 SSA4-GFP::HIS3 MCA1-RFP::LEU2 ssa1Δ::hph ssa2Δ::kanMX4* | This study |
| S1B<br>M1C | *hsp104Δ* guk1-7-GFP | Mat **a** *his3*::pRS403-guk1-7-GFP*::HIS3 leu2Δ0 met15Δ0 ura3Δ hsp104Δ*::*URA3* | This study |
| S1B | *ssa1Δ ssa2Δ hsp104Δ* guk1-7-GFP | Mat **a** *his3*::pRS403-*guk1-7-EGFP::HIS3 leu2Δ0 met15Δ0 ura3Δ ssa1Δ::hph ssa2Δ::kanMX4 hsp104Δ*::*URA3* | This study |
| S1B | *ssa1Δ ssa2Δ hsp104Δ* $P_{GPD}$*SSA4* guk1-7-GFP | Mat **a** *his3*::pRS403-*guk1-7-EGFP leu2Δ0 met15Δ0 ura3Δ ssa1Δ::hph ssa2Δ::kanMX4 hsp104Δ*::*URA3 natMX6:*$P_{GPD}$*-SSA4* | This study |
| S1E | Ssa4-GFP | Mat **a** *his3Δ1 leu2Δ0 met15Δ0 ura3Δ SSA4-EGFP::HIS3* | (Huh et al., 2003) |
| S1E | *ssa1Δ ssa2Δ* Ssa4-GFP | Mat **a** *his3Δ1 leu2Δ0 met15Δ0 ura3Δ ssa1Δ::hph ssa2Δ::kanMX4 SSA4-EGFP::HIS3* | This study |

**TABLE S2: Plasmid list**

| Figures | Name | Replicon | Promoter | Gene | Selection | Source |
|---|---|---|---|---|---|---|
| 3B, C<br>4B, C<br>S2A-D | pGFP-Hsp104 | CEN | *GAL1* | *EGFP-HSP104* | *HIS3* | (Tkach and Glover, 2004) |
| 1B, C, F<br>3D, E<br>5<br>S1D<br>M1 | pguk1-7GFP | - | GPD | *guk1-7-EGFP* | *HIS3* | Per Widlund |
| 1B, C | pgus1-3GFP | - | GPD | *gus1-3-EGFP* | *HIS3* | Per Widlund |
| 2A | pΔssCL* | CEN | Endogenous | *ΔssCPY*-LEU2-myc* | *URA3* | (Eisele and Wolf, 2008) |
| 2B, C | pΔssCG* | CEN | Endogenous | *ΔssCPY*-EGFP-myc* | *URA3* | (Medicherla et al., 2004) |
| 4B, C<br>S2A-D | pRS415 | CEN | GPD | - | *LEU2* | (Sikorski and Hieter, 1989) |
| 4B, C<br>S2A-D | p*SSA1* | CEN | GPD | *SSA1* | *LEU2* | This study |
| 4B, C<br>S2A-D | p*SSA4* | CEN | GPD | *SSA4* | *LEU2* | This study |
| 4B-D<br>S2A-D | p*NBDsw* | CEN | GPD | $NBD_{SSA1}ssa4$ | *LEU2* | This study |
| 4B-D<br>S2A-D | p*SBDsw* | CEN | GPD | $ssa4SBD_{SSA1}$ | *LEU2* | This study |
| 4B, C<br>S2A-D | p*CTDsw* | CEN | GPD | $ssa4CTD_{SSA1}$ | *LEU2* | This study |
| 5 | pSik1-RFP | CEN | GPD | *SIK1-mRFP* | *URA3* | (Sung and Huh, 2007) |
